# Moderate early-life stress improves adult zebrafish (*Danio rerio*) spatial short-term memory but does not affect social and anxiety-like responses

**DOI:** 10.1101/2020.03.10.985945

**Authors:** Barbara D. Fontana, Alistair J. Gibbon, Madeleine Cleal, Ari Sudwarts, David Pritchett, Maria E. Miletto Petrazzini, Caroline H. Brennan, Matthew O. Parker

**Affiliations:** Brain and Behaviour Laboratory, School of Pharmacy and Biomedical Sciences, University of Portsmouth, UK; School of Biological and Chemical Sciences, Queen Mary University London, UK; The International Zebrafish Neuroscience Research Consortium (ZNRC), 309 Palmer Court, Slidell, LA 70458, USA

**Keywords:** Adaptive Flexibility, Moderate-stress, Novel tank, Resilience, Shoal test, FMP Y-maze

## Abstract

Early-life stress (ELS) is defined as a short or chronic period of trauma, environmental or social deprivation, which can affect different neurochemical and behavioral patterns during adulthood. Zebrafish (*Danio rerio*) have been widely used as a model system to understand human neurodevelopmental disorders and display translationally relevant behavioral and stress-regulating systems. In this study, we aimed to investigate the effects of moderate ELS by exposing young animals (six weeks post-fertilization), for three consecutive days, to three stressors, and analyzing the impact of this on adult zebrafish behavior (sixteen weeks post-fertilization). The ELS impact in adults was assessed though analysis of performance on tests of unconditioned memory (free movement pattern Y-maze test), exploratory and anxiety-related task (novel tank diving test) and social cohesion (shoaling test). Here, we show for the first time that moderate ELS increases the number of pure alternations compared to pure repetitions in the unconditioned Y-maze task, suggesting increased spatial short-term memory, but has no effect on shoal cohesion, locomotor profile or anxiety-like behavior. Overall, our data suggest that moderate ELS may be linked to adaptive flexibility which contributes to build ‘resilience’ in adult zebrafish by improving short-term spatial memory performance.

## 1. Introduction

Early□life experiences are often linked to changes in cognitive and behavioral aspects in humans (Pechtel & Pizzagalli, 2011). In animal models, early-life stress (ELS) is shown to directly cause long-term changes in several brain functions (Molet, Maras, Avishai-Eliner, & Baram, 2014; Nishi, Horii-Hayashi, Sasagawa, & Matsunaga, 2013; Spinelli et al., 2009). ELS can occur prenatally or postnatally (or both), and can affect both neurological and physiological development (Maniam, Antoniadis, & Morris, 2014; Pechtel & Pizzagalli, 2011). The neurochemical and hormonal changes induced by ELS are associated with emotional and cognitive processes, and ELS is a risk factor for neuropsychiatric disturbances such as depression (Coffino, 2009; Heim & Nemeroff, 2001; Kaufman, Plotsky, Nemeroff, & Charney, 2000), posttraumatic-stress disorder (PTSD), schizophrenia, substance abuse (Scheller-Gilkey, Moynes, Cooper, Kant, & Miller, 2004), attention-deficit disorder (Heim & Nemeroff, 2001), eating disorders (Gilbert et al., 2009), and an abnormal stress response (Kajantie & Raikkonen, 2010; Kaufman et al., 2000) in adult life.

ELS can be assessed behaviorally and biologically in rodents by using protocols such as maternal separation, post-weaning social isolation and peripubertal stress (Dunphy-Doherty et al., 2018; Lyons, Parker, & Schatzberg, 2010; Ouchi et al., 2018; Tsuda, Yamaguchi, & Ogawa, 2011). Although high or chronic levels of stress may disturb brain development and affect behavior, acute activation of the body’s stress response systems can adapt and increase chances of survival (Anda et al., 2006; De Bellis et al., 1999; Lupien, McEwen, Gunnar, & Heim, 2009; Maniglio, 2009; Pechtel & Pizzagalli, 2011; Spataro, Mullen, Burgess, Wells, & Moss, 2004). For example, *predictable* chronic stress increases ‘resilience’ (i.e. positive outcomes in spite of adversity environmental or experiences) in rats by increasing hippocampal neurogenesis and memory (Parihar, Hattiangady, Kuruba, Shuai, & Shetty, 2011), whereas *unpredictable* chronic stress may impair learning and memory processes (Rice, Sandman, Lenjavi, & Baram, 2008). Moreover, ELS can change other behavioral domains such as stress reactivity (O’Mahony et al., 2009) and impulsivity (Lovallo, 2013). In general, because neuropsychiatric studies depict only deleterious outcomes following ELS (Grassi-Oliveira, Ashy, & Stein, 2008; Teicher et al., 2003; Teicher, Tomoda, & Andersen, 2006; Weber & Reynolds, 2004), relatively little attention has been devoted to evaluating positive cognitive outcomes of ELS. Thus, a better understanding of the long-term consequences of ELS could improve prevention strategies for mental disorders, particularly for those that are affected by stress.

Zebrafish (*Danio rerio*) are widely utilized as a translational and complementary model to better understand human neurodevelopmental-related disorders (Sakai, Ijaz, & Hoffman, 2018). This usage is primarily due to high genetic (70% of zebrafish genes have at least one human orthologue) (Howe et al., 2013) and physiological homology (Holzschuh, Ryu, Aberger, & Driever, 2001; MacRae & Peterson, 2015; Rico et al., 2011) between fish and other vertebrates. Although there are several differences between human and zebrafish stress-regulating systems, there is conservation of organization and functioning in terms of anatomy, connectivity, and molecular constituents, making zebrafish an important model to study stress-related responses (D. Alsop & M. Vijayan, 2009; Alsop & Vijayan, 2008; D. Alsop & M. M. Vijayan, 2009; Grunwald & Eisen, 2002). Zebrafish display a broadly conserved behavioral repertoire (Kalueff et al., 2013), including different domains such as learning (Cognato Gde et al., 2012; Valente, Huang, Portugues, & Engert, 2012), aggressiveness (Ariyomo, Carter, & Watt, 2013; Norton, 2018), anxiety-like behavior (Blaser & Rosemberg, 2012) and social behavior (de Polavieja & Orger, 2018; Dreosti, Lopes, Kampff, & Wilson, 2015; Paull et al., 2010). Like mammals, zebrafish respond to a variety of stressors such as handling, social isolation, rapid temperature changes, overcrowding and novel environments, operationally defined through increases in abnormal behaviors such as decreased locomotion and social interactions (Fulcher, Tran, Shams, Chatterjee, & Gerlai, 2017; Pavlidis et al., 2013; Piato et al., 2011). Overall, zebrafish have great potential as a translational model for studying ELS. Here, we investigated the effects of moderate ELS by exposing young animals (six weeks post-fertilization) for three consecutive days, to three stressors, and analyzing the impact of this on adult zebrafish behavior (sixteen weeks post-fertilization) through analysis of performance on tests of unconditioned memory (free movement pattern (FMP) Y-maze test), exploratory and anxiety-related task (novel tank diving test) and social cohesion (shoaling test).

## 2. Materials and Methods

### 2.1. Animals, blinding and randomization

Experiments were carried out using Tubingen (TU) zebrafish bred in house **following breeding from multiple pairs of fish**. Fish were sorted into groups of 30-40, and grown to 6-weeks post fertilization in a recirculating nursery system (Tecniplast). At this point, the ELS protocol was carried out (see below for details). All fish were then grown to four-months post fertilization in groups of ∼20, and were housed in groups according to stress protocol (ELS vs handling control; 5 x groups of each condition). The experimental and technical teams were blind as to treatment allocation during rearing. Adult (4-months post fertilization) animals were tested on one of the three behavioral procedures to ascertain perseveration/spatial short-term memory (FMP Y-maze test), exploratory and anxiety-related task (novel tank diving test) and sociability (shoaling test). Animals were maintained on a 14/10-hour light/dark cycle (lights on at 9:00 a.m.), pH 7.1, at ~28.5 °C (±1 °C) and were fed three times/day with a mixture of live brine shrimp and flake food, except during the weekend, when they were fed once/day. Each animal was tested in only one behavioral protocol to avoid potential effects of multiple testing. Animals were euthanized by using 2-phenoxyethanol (Aqua-Sed™, Vetark, Winchester, UK) and all experiments were carried out following scrutiny by the University of Portsmouth Animal Welfare and Ethical Review Board and under license from the UK Home Office (Animals (Scientific Procedures) Act, 1986) [PPL: P9D87106F]. All stressors and behavioral tasks were carried out in a fully blinded manner and once all data were collected and screened for extreme outliers (e.g., fish freezing and returning values of ‘0’ for behavioral parameters indicating non-engagement), group allocation (control or ELS) was revealed and data analyzed in full. Final sample sizes for all behavioral tests were established following power analyses calculated with pilot experiments/prior publications from our group.

### 2.2. Short-term ELS protocol

The stressors were applied each day for three consecutive days. During this time, the fish were exposed to one of three stressors on each day. These included ‘water change’ (water from tank house changed for 3 times), ‘shallow water’ (2 min in shallow water; **60% of body exposure to the air**), and ‘overcrowding’ (10 fish/well in a 6 well-plate for 45 min) (**Fig. 1A**). The three stressors used here were adapted from previous studies where water change, shallow water and overcrowding induced behavioral and cortisol changes (Clark, Boczek, & Ekker, 2011; Ramsay et al., 2006). Furthermore, all the three stressors were separately tested to investigate if they were able to modulate cortisol levels in animals with 6 weeks old. The randomization schedule for exposure to stressors is displayed in **Table 1**.

**Table 1.** Procedure of the mild stress protocol in larvae zebrafish

| Group | N | Day 1 | Day 2 | Day 3 | 621 |
| --- | --- | --- | --- | --- | --- |
| S1 | 25 | O,S,W | S,O,W | W,O,S |  |
| S2 | 25 | W,O,S | O,S,W | S,O,W | 622 |
| S3 | 26 | S,O,W | W,O,S | O,S,W |  |
| C1 | 25 | - | - | - | 623 |
| C2 | 25 | - | - | - |  |
| C3 | 25 | - | - | - | 624 |

625 **Abbreviations:** Overcrowding (O); Shallow Water (S); Water change (W).

**Figure 1.**
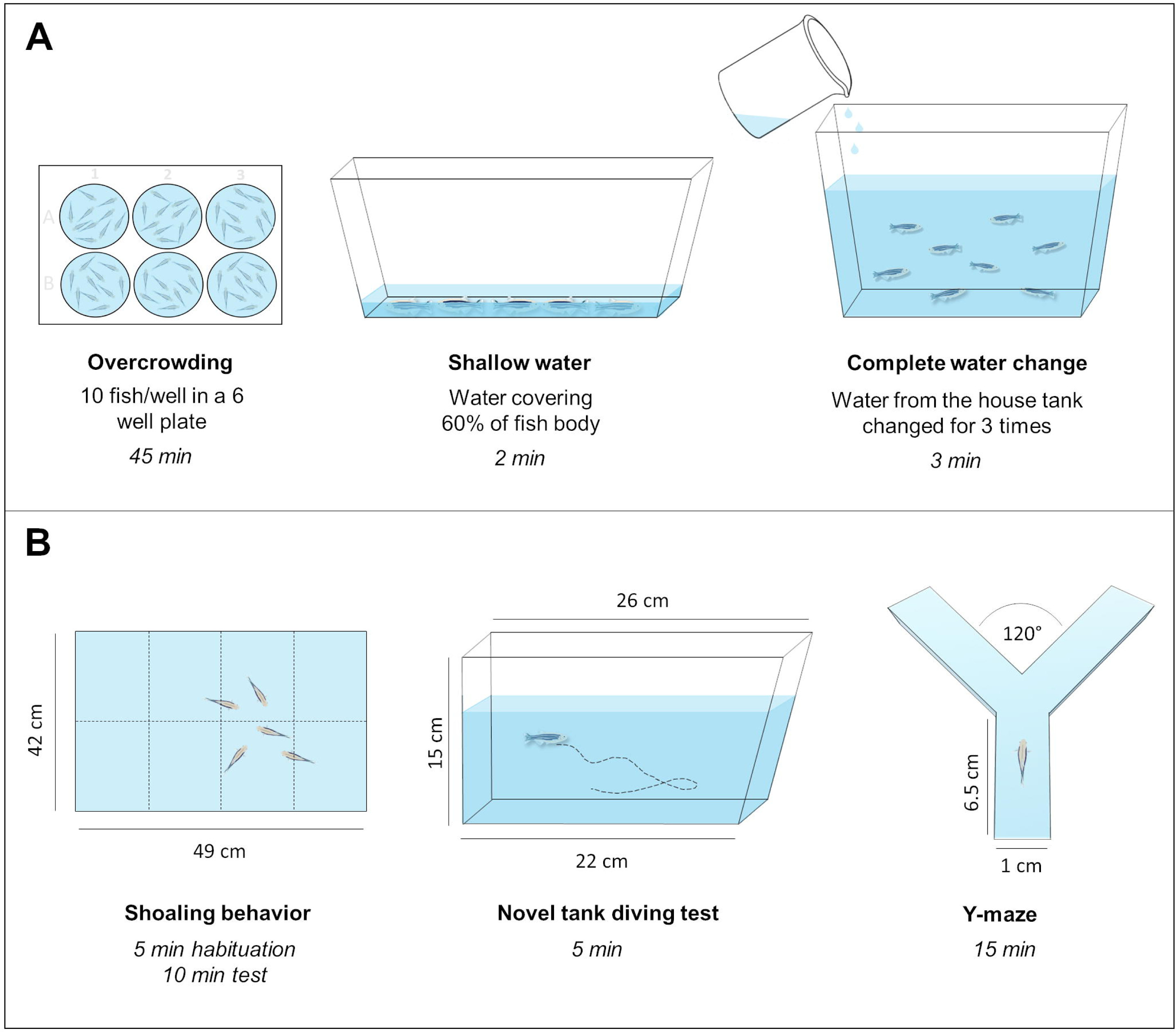
Schematic representation of the **(A)** stressors and **(B)** behavioral tasks used to assess ELS in adult zebrafish.

### 2.3. Novel tank diving test

The novel tank diving test is commonly used for analyzing exploratory and locomotor profiles. Moreover this task is often used to analyze anxiety-like behavior being sensitive to a range of anxiogenic and anxiolytic drugs in absence or presence of different aversive stimulus (Egan et al., 2009; Maximino et al., 2018; Mezzomo, Silveira, Giuliani, Quadros, & Rosemberg, 2016; Wong et al., 2010). Zebrafish (*n= 23*) were placed individually in a novel tank (22 - 26 cm length x 15 cm height x 9 cm width) (**Fig. 1B**). Behavioral activity was recorded in front of the tank using a webcam for 5 min to analyze diving response (Egan et al., 2009; Parker et al., 2012; Rosemberg et al., 2012). The tank was virtually divided in three areas (bottom, middle and top) and time at the bottom was used to assess anxiety-like phenotypes. The total distance traveled was used to evaluate differences in locomotor profile. All behaviors were measured with automated video-tracking software (EthoVision, Noldus Information Technology Inc., Leesburg, VA - USA) at a rate of 60 frames/s.

### 2.4. Shoaling test

Zebrafish is a social species that presents a natural preference for conspecifics in neutral and mildly stressful situations (de Polavieja & Orger, 2018; Paull et al., 2010; Saverino & Gerlai, 2008). Social cohesion (*n=80*, organized into 16 groups of *n= 5*) was tested in a shoaling assay. Sixteen groups of n = 5 fish from each treatment (ELS vs no stress) were placed in a white opaque tank (49 cm length x 15 cm height x 42 cm width) filled with 6L of aquarium treated water (**Fig. 1B**). Animals were left to habituate for 5 mins, then filmed for 10-min from above by using a webcam. The unit of replication was shoal (i.e., group of 5 fish) nested in housing group (*n =8* shoals per treatment). Data analysis was based on previous work (Parker et al., 2014; Parker, Brock, Millington, & Brennan, 2013). Briefly, the arena was split into 8 equal sections and the number of fishes in each square (as a function of total number of squares occupied) at any one time was ascertained every 10-seconds to calculate a dispersion index. If a fish was between squares, the square containing >50% of the fish was recorded as the occupied square. Shoaling behavior was measured by using the automated video-tracking software (EthoVision, Noldus Information Technology Inc., Leesburg, VA - USA) at a rate of 60 frames/s.

### 2.5. FMP Y-maze

The FMP Y-maze is used for measuring spatial short-term memory in rodents and zebrafish. Spontaneous alternations are reduced following administration of memory-impairing drugs (eg MK801 and scopolamine), suggesting that alternations are indicative of spatial short-term memory (Barnard, Matthews, Messori, Podaliri-Vulpiani, & Ferri, 2016; Castellano, Diaz-Palarea, Rodriguez, & Barroso, 1987; Cunha et al., 2008; Fontana, Cleal, Clay, & Parker, 2019; Frasnelli, 2013; Rodriguez & Afonso, 1993; Rodriguez, Gomez, Alonso, & Afonso, 1992). Fish (*n=48*) were placed, individually, in a Y-shaped maze (6.5 cm length x 1 cm width; three identical arms at a 120° angle from each other) with white opaque walls (**Fig. 1B**), and recorded for 15 minutes. During this time, the fish could either turn left or right at each choice point. Ambient light allowed some visibility in the maze, but no explicit intra-maze cues were added to the environment. All responses were non-reinforced. Turn choices were organized in blocks of four trials, and within each 4-trial block, the fish could either show pure alternation (rlrl, lrlr) or pure repetition (llll, rrrr), or any combination between (16 in total). Therefore, FMP Y-maze behavior was analyzed through 16 possible four-trial outcomes (tetragrams) for each fish as a proportion of the total number of turns to analyze, in detail, their behavior patterns (Cleal & Parker, 2018; Gross, Engel, Richter, Garner, & Wurbel, 2011). If behavior was fully random, it would be predicted that each choice would be equally likely (i.e., the predicted frequencies would be equal for each tetragram). However, if behavior was biased towards alternation, or repetition, it would be predicted that the alternation and repetition tetragrams would be significantly different from one another.

### 2.5. Cortisol levels

For the analysis of cortisol levels, fish (6 weeks old) were acutely exposed to water change, shallow water or overcrowding using the conditions previously described. Due to their small size, animals were pooled in *n = 12* per sample. Cortisol levels were assessed using a human salivary cortisol ELISA kit (Salimetrics) as previously described (Cachat et al., 2010; Parker, Millington, Combe, & Brennan, 2012). Fish were killed by immersion in ice-water and the whole bodies were **snap-frozen** in liquid nitrogen then frozen at −80C until assay. Samples were homogenized in 5 ml ice-cold PBS. 5 ml of diethyl ether was added, and samples were centrifuged (7000 x g) for 15 minutes, and the top (organic) layer was removed. This process was repeated three times and then the diethyl ether was evaporated overnight. The resulting cortisol was reconstituted in 1 ml ice cold PBS and the ELISA was then performed in 96-well plates as per the manufacturer’s instructions. Cortisol concentrations (ng/g-1) were determined from OD readings compared against manufacture standards. All samples were run in duplicate and the inter- and intra-assay coefficients of variation were %.

### 2.6. Statistics

Data were analyzed in IBM SPSS® Statistics and results were expressed as means ± standard error of the mean (S.E.M). To assess whether there were any effects of stress on total turns, pure alternations (lrlr + rlrl), pure repetitions (rrrr + llll), right turns, left turns, distance traveled, time at the bottom and shoaling ratio, we used Student’s T-Tests. Additionally, to investigate the stress effects on behavioral tetragrams we fitted a generalized linear mixed effects model (Poisson distribution, log link), with stress and block choice as fixed factors, and ID as a random effect (to account for non-independence of replicates). Cortisol levels were analyzed by using one-way ANOVA. When appropriate, Tukey’s test was used as post-hoc analysis, and results were considered significant when p ≤ 0.05.

## 3. Results

### 3.1. Cortisol levels are increased in stressed young animals

Initially, we exposed small groups of 6-week old fish to the three stressors (shallow water, water change, and overcrowding) and examined the cortisol response to the exposures. A one-way ANOVA confirmed that the treatments were all invoking a stress response, with a significant difference between groups for cortisol levels (F _(3, 8)_ = 10.67; *p* = 0.0036). Post-hoc tests revealed that cortisol levels were significantly increased when acutely exposing 6 weeks animals to each of the three stressors as compared to no treatment controls: control < shallow water (*p<* 0.005), control < water change (*p<* 0.05) and control < overcrowding (*p<* 0.05) (Fig 2). All samples were run in duplicate and the inter- and intra-assay coefficients of variation were <4 %.

**Figure 2.**
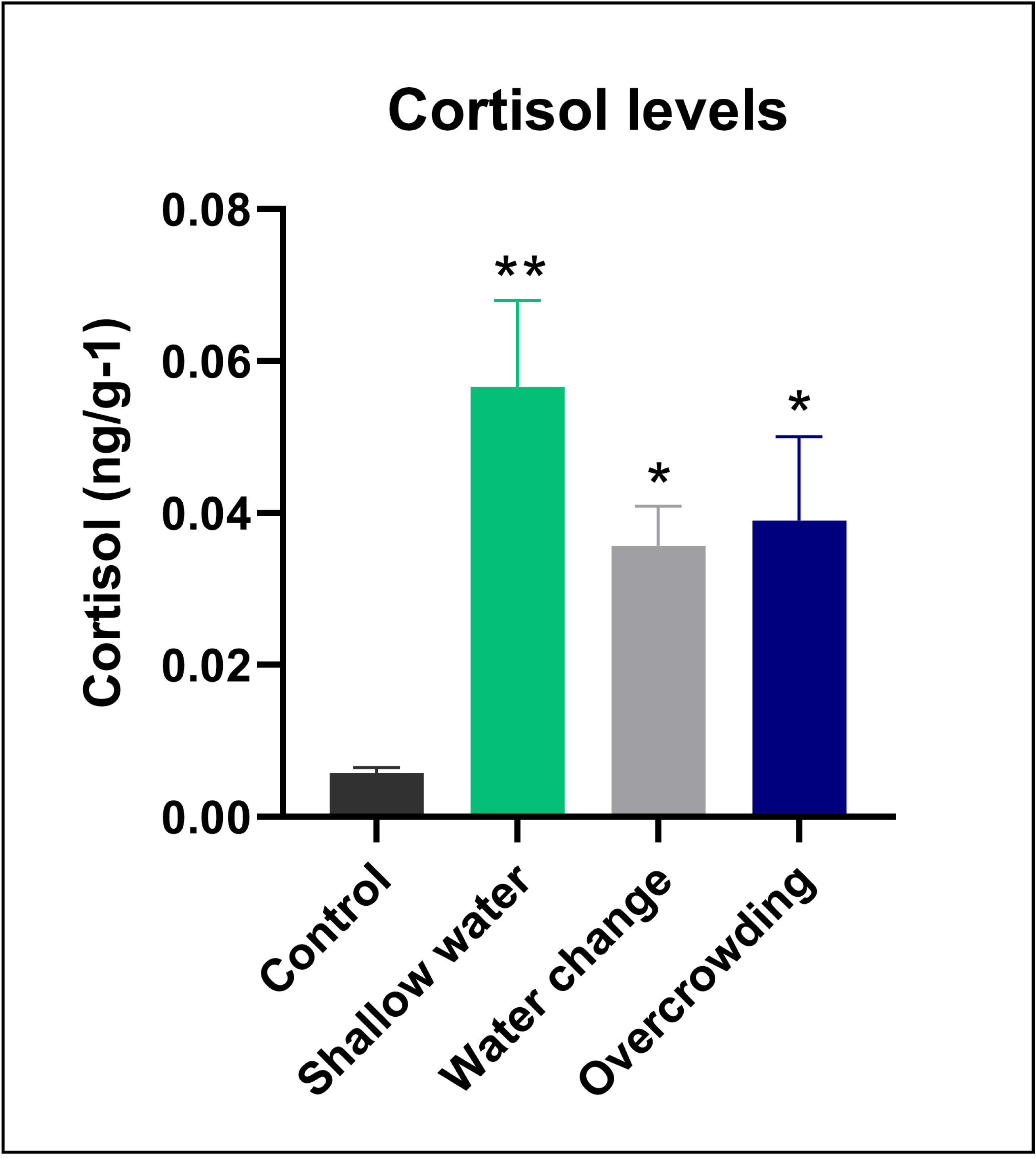
Cortisol levels for non-stressed (control) and stressed animals. Data were represented as mean ± S.E.M and analyzed by one-way ANOVA, followed by Tukey’s test multiple comparison test. Asterisks indicates statistical differences compared to control group (*p < 0.05 and **p <0.01, *n* = 3 −4 *per* group).

### 3.2. Anxiety and social-related phenotypes are not modulated by short-term ELS

We examined the effects of ELS on anxiety using the novel tank test, and social behavior using a social cohesion assay. Student’s t-tests (two-tailed) were performed for the analyses of anxiety-like behavior and shoal responses. We observed that ELS did not change any locomotor nor anxiety-related patterns such as distance travelled (t_(54)_ = 0.868; *p* = 0.388) and time spent in the bottom section (t_(54)_ = 0.166; p = 0.86) (**Fig. 3A**). There was also no difference in the social cohesion ratio (t_(54)_ = 0.060; *p* = 0.952) observed for animals exposed to ELS compared to the control group (**Fig. 3B**).

**Figure 3.**
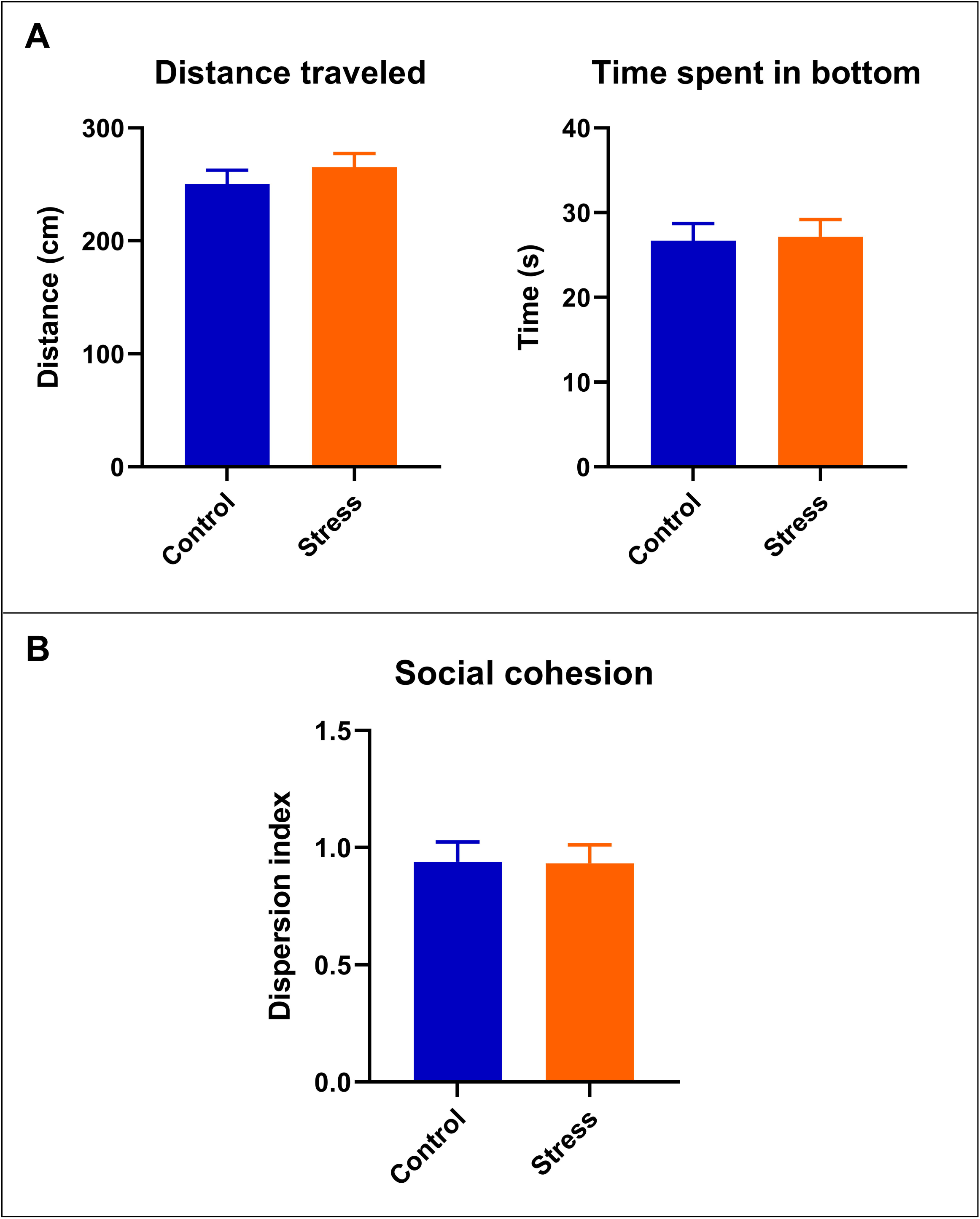
Effects of ELS in stress-reactivity and social responses. **(A)** ELS do not change distance traveled and time spent in bottom in the novel tank diving task (*n* = 11 – 12 *per* group). **(B)** Shoal cohesion is not affected by three-days of ELS in adult zebrafish (*n* = 8 *per* group). Data were represented as mean ± S.E.M. and analyzed by Student’s T-Test.

### 3.3. Short-term spatial memory is enhanced in adult zebrafish exposed to ELS

Finally, we examined the effects of ELS on short term spatial memory by examining turn choices in the FMP Y-maze exploration task. Through analysis of overall turn choices, we found that ELS affected FMP Y-maze performance. Fig. 4 depicts the behavioral 16 choice tetragrams for control *vs* ELS exposed animals. There was no main effect of ELS on tetragram analysis (F _(1, 336)_ = 0.00; *p* > 0.999). There was, however, a main effect of block choice (F _(15, 336)_ = 54.75; *p* < 0.0001), and a treatment * block-choice interaction (F _(15, 336)_ = 8.724; *p* < 0.0001). Post-hoc Tukey tests revealed that ELS significantly increased the rlrl (*p*< 0.0001) and lrlr (*p*< 0.0001) block choices (pure alternations). Examining alternations and repetitions in more detail revealed that ELS significantly decreased the number of total turns (t_(21)_ = 2.846; *p* = 0.009), and number of repetitions (t_(21)_ = 3.055; *p* = 0.006) while increasing the number of alternations (t_(21)_ = 3.055; *p* = 0.0012) in the FMP Y-maze test (Fig. 5). ELS did not alter animals preference for left (t_(21)_ = 0.533; *p* = 0.599) or right turns (t_(21)_ = 0.533; *p* = 0.599).

**Figure 4.**
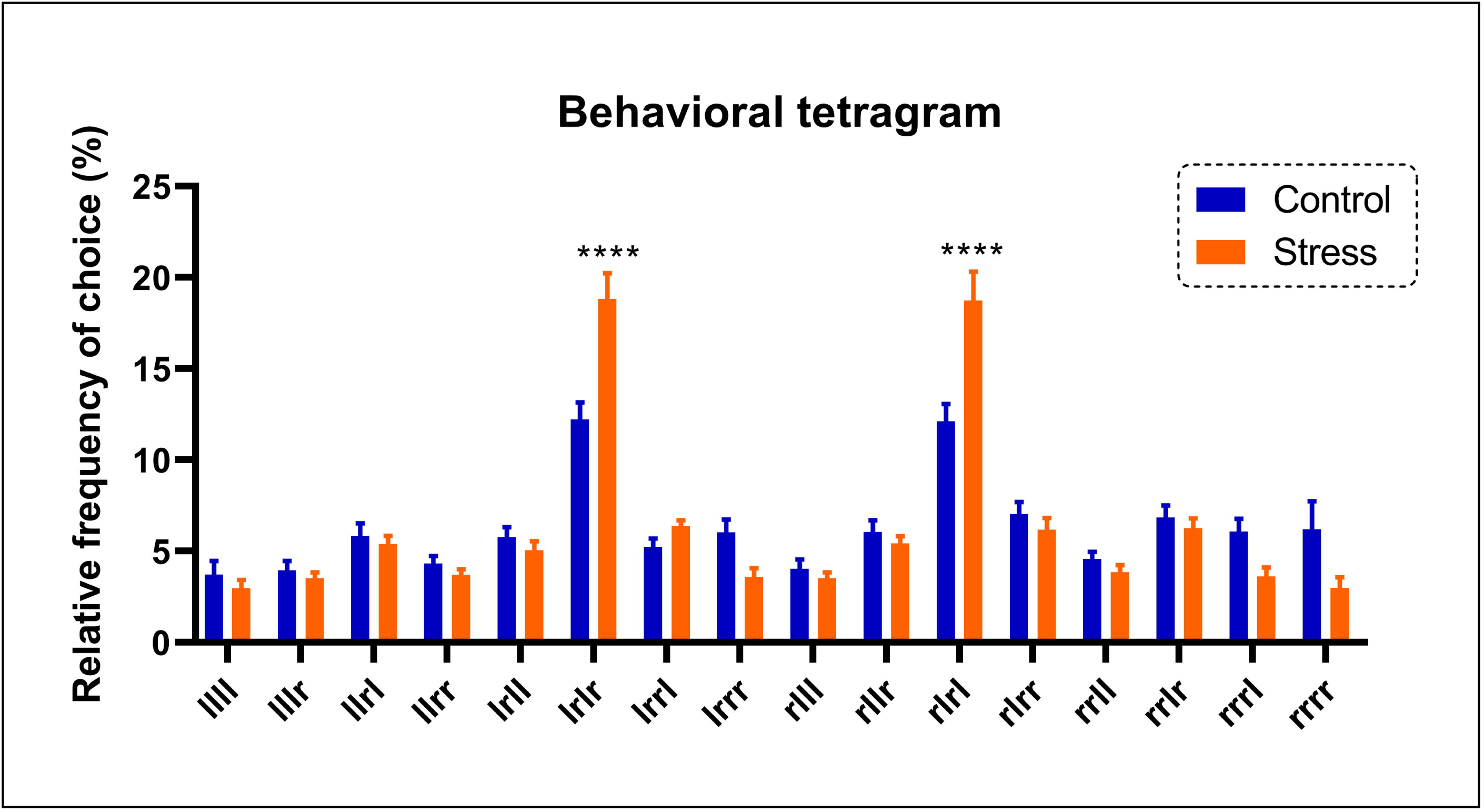
Y-maze tetragrams showing the behavioral phenotype of control and ELS groups. Data were represented as mean ± S.E.M. and analyzed by linear mixed effects, followed by Tukey’s multiple comparison test. Asterisks indicates statistical differences compared to non-biased group or between biased groups (*p < 0.05, **p <0.01, ***p <0.001 and *p<0.0001, *n* = 24 *per* group).

**Figure 5.**
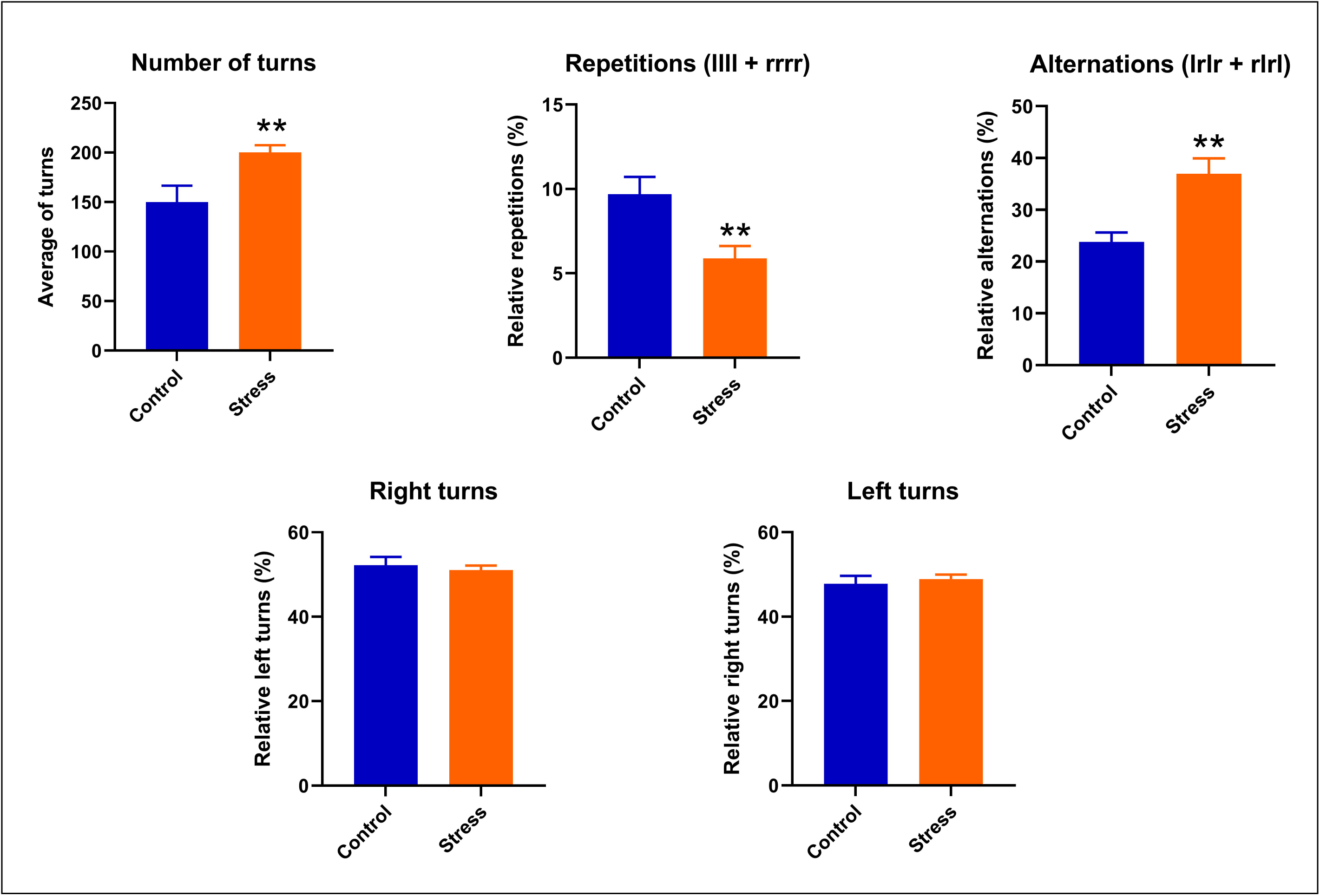
ELS increase alternations and decrease repetition without change left or right preference of adult zebrafish. Data were represented as mean ± S.E.M. and analyzed by Student’s T-Test. Asterisks indicates statistical differences compared to control group (*p < 0.05 and **p <0.01, *n* = 24 *per* group).

## 4. Discussion

In this study we evaluated the effects of exposing young zebrafish (16 days old) to unpredictable moderate stress for three days on adult behavior (four months old), through the analysis of three different domains: locomotor and anxiety (novel tank diving test), sociality (shoal cohesion test) and memory (FMP Y-maze test). We showed, for the first time, that three days of ELS improves zebrafish short term spatial memory by increasing pure alternations and decreasing repetitions in the FMP Y-maze test. Although learning and memory process were affected by short-term ELS, stress did not modulate social behavior and anxiety-like behavior in adult zebrafish. Collectively, these data support previous evidence that early-life exposure to moderate stress can build resilience and have an important adaptive role for different species, including humans. Our study has highlighted that zebrafish is an important animal model for elucidating the developmental processes underlying ELS, and how this impacts adult behavior.

Early-life experiences regulate individual differences through both neural plasticity and epigenetic modifications, and these modifications later determine the capacity for flexible and adaptive behaviors in adulthood (McEwen, Bowles, et al., 2015; McEwen, Gray, & Nasca, 2015). Stressful stimuli can exert both positive and negative impacts on neurophysiological aspects being balanced to give the best fitness outcome (Bogdan & Pizzagalli, 2006; Gluckman, Hanson, & Beedle, 2007; Maier, Amat, Baratta, Paul, & Watkins, 2006; Zannas & West, 2014). To confirm if shallow water, water changes and overcrowding changed zebrafish stress-regulating systems, we analyzed the effects of these three stressors separately in animals with 6 weeks old. As previously observed in adults (Pavlidis et al., 2013; Song et al., 2018), the acute stressors increased cortisol levels in young animals.

The impact on adult life of ELS follows an inverted U-shaped curve, where exposure to mild/moderate stress during development results in positive/beneficial outcomes such as resilience, and either too little or too much stress exposure (or too high an intensity) leads to negative outcomes (Russo, Murrough, Han, Charney, & Nestler, 2012; Sapolsky, 2015). Concerning this, chronic ELS is related to different behavioral outcomes in different species, including enhanced emotional- and stress-reactivity, which is frequently linked to anxiety disorders (Coplan et al., 1996; Nugent, Tyrka, Carpenter, & Price, 2011), aggressiveness (Veenema, 2009; Veenema, Blume, Niederle, Buwalda, & Neumann, 2006), impulsiveness (Lovallo, 2013) and antisocial behavior (Haller, Harold, Sandi, & Neumann, 2014; Kohl et al., 2015). Meanwhile, adolescents exposed to moderately severe ELS events showed blunted depressive symptom responses to changes in proximal stressful events in the previous 9 months, compared to those with fewer ELS events (Shapero et al., 2015). Here, we showed that short-term ELS did not affect adult zebrafish anxiety-like profile nor social cohesion in both the novel tank diving test and shoal cohesion test. However, ELS increased the number of pure alternations, and decreased the number of pure repetitions, in the FMP Y-maze task.

The FMP Y-maze task comprises three identical arms, where the animal is introduced to the center of the maze and allowed to freely explore. In this task, over the course of multiple arm entries, animals have a natural tendency to enter the recently visited arm less frequently, thus increasing the number of alternations across the test (Drew, Miller, & Baugh, 1973; Hughes, 2004; Kokkinidis & Anisman, 1976; Swonger & Rech, 1972). In general, various configurations of the Y-maze task have been used to study learning and processes in animal models, including rodents (Fu et al., 2017; Ghafouri et al., 2016; Luine, 2015) and fish (Aoki, Tsuboi, & Okamoto, 2015; Cleal & Parker, 2018; Cognato Gde et al., 2012). Both decreased alternations and increased repetitions are affected pharmacologically by muscarinic and NMDA-receptor antagonists, and additionally by β-amyloid peptides in rodents (Cunha et al., 2008; Hiramatsu & Inoue, 2000; Park et al., 2010; Walker & Gold, 1992). Recently, we also observed that zebrafish presented decreased alternations when exposed to moderate alcohol during early nervous system developmental (Cleal & Parker, 2018). Although the FMP Y-maze responses can be influenced by novelty-seeking (Fontana et al., 2019), we observed here that ELS did not change exploratory responses of animals at the novel tank diving task. Thus, it seems likely, that increased pure alternations and a coincidental decrease in pure repetitions are indicative that the ELS animals may have an improvement in learning and memory adaptive flexibility, thus suggesting increased ‘resilience’ as a result of the ELS exposure.

Memory is a highly dynamic process that is built from initial encoding, to the new and fragile memory trace that is stabilized during consolidation and reactivated during memory retrieval (Abel & Lattal, 2001; Bisaz, Travaglia, & Alberini, 2014). Memory can also return to an unstable state in which reconsolidation is needed to stabilize it (Dudai, 2006; Lv, 2015). Stress has been shown to affect all phases of memory (encoding, consolidation, memory retrieval and reconsolidation); however, how stress influences memory depends on *when* an individual is stressed, and what *frequency* and *intensity* of that stress are (Schwabe & Wolf, 2012). When looking at resilience induced by stress, it seems that recovery from stress-inducing changes in neural architecture is not simply a reversal, but instead is a form of neuroplastic adaptation (McEwen, Gray, et al., 2015). Although neural plasticity and epigenetic factors are known to underlie the mechanisms responsible for ELS inducing resilience in adults (Cadet, 2016; Kentner, Cryan, & Brummelte, 2018; McEwen, Gray, et al., 2015), there remains no clear understanding of the mechanisms involved in these processes and how they are functionally associated.

## 5. Conclusion

Here, we show for the first time that adult zebrafish exposed to moderate ELS can build resilience as evidenced by increases in spatial short-term memory, but without changing anxiety-like behavior or social behavior patterns. We suggest that this protocol could serve as an useful model to understand: a) translational genetic and physiological aspects are associated with memory adaptive flexibility; b) the evolutionary and conserved characteristics that are common between zebrafish and humans that correspond to stress-induced changes in memory and learning. Overall, future studies should investigate how moderate ELS can modulate neuroplastic adaptation and epigenetics in zebrafish adaptive flexibility and resilience.

## Acknowledgments

This study was financed in part by the Coordenação de Aperfeiçoamento de Pessoal de Nível Superior - Brazil (CAPES) - Finance Code 001 at the University of Portsmouth, UK (BDF). MC is supported by a Science Faculty Studentship from the University of Portsmouth. CHB is supported by BBSRC (BB/M007863/1), leverhulme grant (RPG-2016-143), Human Frontiers grant (HFSP - RGP0008/2017).

