## Supplementary material for "Moderate early-life stress improves adult zebrafish (*Danio rerio*) spatial short-term memory but does not affect social and anxiety-like responses": Table 1

| Group | N | Day 1 | Day 2 | Day 3 |
| --- | --- | --- | --- | --- |
| S1 | 25 | O,S,W | S,O,W | W,O,S |
| S2 | 25 | W,O,S | O,S,W | S,O,W |
| S3 | 26 | S,O,W | W,O,S | O,S,W |
| C1 | 25 | - | - | - |
| C2 | 25 | - | - | - |
| C3 | 25 | - | - | - |

**Table 1.** Procedure of the mild stress protocol in larvae zebrafish

**Abbreviations:** Overcrowding (O); Shallow Water (S); Water change (W).
